# VISTA-induced tumor suppression by a four amino acid intracellular motif

**DOI:** 10.1101/2025.01.05.631401

**Authors:** Yan Zhao, Tina Andoh, Fatima Charles, Priyanka Reddy, Kristina Paul, Harsh Goar, Ishrat Durdana, Caiden Golder, Ashley Hardy, Marisa M. Juntilla, Soo-Ryum Yang, Chien-Yu Lin, Idit Sagiv-Barfi, Benjamin S. Geller, Stephen Moore, Dipti Thakkar, Jerome D. Boyd-Kirkup, Yan Peng, James M. Ford, Melinda L. Telli, Song Zhang, Allison W. Kurian, Robert B. West, Tao Yue, Andrew M. Lipchik, Michael P. Snyder, Joshua J. Gruber

**Author notes:** Corresponding Authors: Joshua Gruber, Michael Snyder.

## Abstract

VISTA is a key immune checkpoint receptor under investigation for cancer immunotherapy; however, its signaling mechanisms remain unclear. Here we identify a conserved four amino acid (NPGF) intracellular motif in VISTA that suppresses cell proliferation by constraining cell-intrinsic growth receptor signaling. The NPGF motif binds to the adapter protein NUMB and recruits Rab11 endosomal recycling machinery. We identify and characterize a class of triple-negative breast cancers with high VISTA expression and low proliferative index. In tumor cells with high VISTA levels, the NPGF motif sequesters NUMB at endosomes, which interferes with epidermal growth factor receptor (EGFR) trafficking and signaling to suppress tumor growth. These effects do not require canonical VISTA ligands, nor a functioning immune system. As a consequence of VISTA expression, EGFR receptor remains abnormally phosphorylated and cannot propagate ligand-induced signaling. Mutation of the VISTA NPGF domain reverts VISTA-induced growth suppression in multiple breast cancer mouse models. These results define a mechanism by which VISTA represses NUMB to control malignant epithelial cell growth and signaling. They also define distinct intracellular residues that are critical for VISTA-induced cell-intrinsic signaling that could be exploited to improve immunotherapy.

## Introduction

The V-domain immunoglobulin suppressor of T cell activation (VISTA) checkpoint molecule is an immunoglobulin domain-containing surface receptor with close sequence homology to PD-L1 [1] expressed throughout the hematopoietic system. In T cells, VISTA expression is highest in the naïve compartment and loss of VISTA leads to increased T cell activation and contraction of the naïve pool [2]. Mice lacking VISTA have increased susceptibility to a range of autoimmune or inflammatory conditions including lupus, encephalitis, and atopy [1, 3–10]. In cancer models, VISTA blockade is sufficient to induce antitumor immunity and can potentiate other immunotherapies [11–14]. Therefore, clinical trials are testing VISTA-blocking small molecules and antibodies for human cancer (NCT04475523, NCT02812875, NCT04564417, NCT05082610).

Molecular mechanisms of VISTA biology are still being elucidated. Multiple receptors can interact with the extracellular domain of VISTA, including VSIG3, PSGL1, LRIG1 and VISTA itself [15–18]. However, it remains unclear whether the main action of VISTA is through its extracellular domain binding to a purported ligand or through cytoplasmic signaling events transmitted via VISTA’s intracellular domain. For example, T cells lacking VISTA show increased proliferation and cytokine production in response to T cell receptor (TCR) crosslinking, suggesting a role for VISTA in intracellular signaling [3, 19]. However, treatment of T cells with a recombinant VISTA-Fc fusion disrupted CD3-mediated TCR activation, implying that VISTA may function through binding a cell surface receptor [19]. Additional roles for VISTA in Src activation [20], or signaling through ERK and STAT pathways [21] have been proposed. However, thus far the field lacks a unified mechanism that explains how VISTA controls immunity.

Triple-negative breast cancer (TNBC) is characterized by an aggressive clinical course with rapid cell division, genetic heterogeneity, and poor outcomes [22, 23]. However, there is significant heterogeneity in TNBC proliferative potential, immunological infiltration and spectrum of mutations [24–29]. Treatment relies primarily on cytotoxic chemotherapies, and more recently PD1-targeted immunotherapy [30, 31]. For patients with metastatic TNBC, only ∼40% of patients qualify for existing immunotherapy drugs [32, 33]. This has led to interest in identifying other potential immunotherapeutic pathways for TNBC.

We recently published multi-omic epigenetic profiling of mammary cells grown in epidermal growth factor (EGF), which identified an immunological signature associated with NF-κB activity [34, 35]. Here, we report that VISTA expression in cell lines and human tumors suppresses tumor growth. Importantly, these effects were cell-intrinsic and did not require a functional immune system, nor expression of canonical VISTA ligands. This led us to focus on VISTA’s intracellular domain, where we identify a functional four amino acid motif that is required for VISTA-mediated cell growth suppression in cancer cell lines and tumor xenografts. These results provide the first unified molecular mechanism to explain how VISTA represses cell proliferation by binding adapter proteins to control cell-intrinsic co-receptor signaling.

## Results

### VISTA+ TNBCs have low PD-L1 and diminished proliferative index

Multi-omic analyses including RNA-seq and ATAC-seq were previously used to identify epigenetic signatures induced in mammary cells [34]. We used NF-κB transcription factor-binding motifs to focus on immunoregulator genes (n=3729 genes; **Fig. 1a**). The *VSIR* gene (which encodes the VISTA protein, also known as PD-1H and B7-H5) was identified as more highly expressed in cells grown without EGF compared to EGF-stimulated cells (**Fig. 1b, c**). This effect was specific to VISTA because transcript and protein levels of PD-L1 and PD-L2 were suppressed by EGF withdrawal (**Fig. 1d, e**). Therefore, *VSIR* gene expression was suppressed by growth factor stimulation and induced by growth factor withdrawal in mammary cell lines.

**Figure 1:**
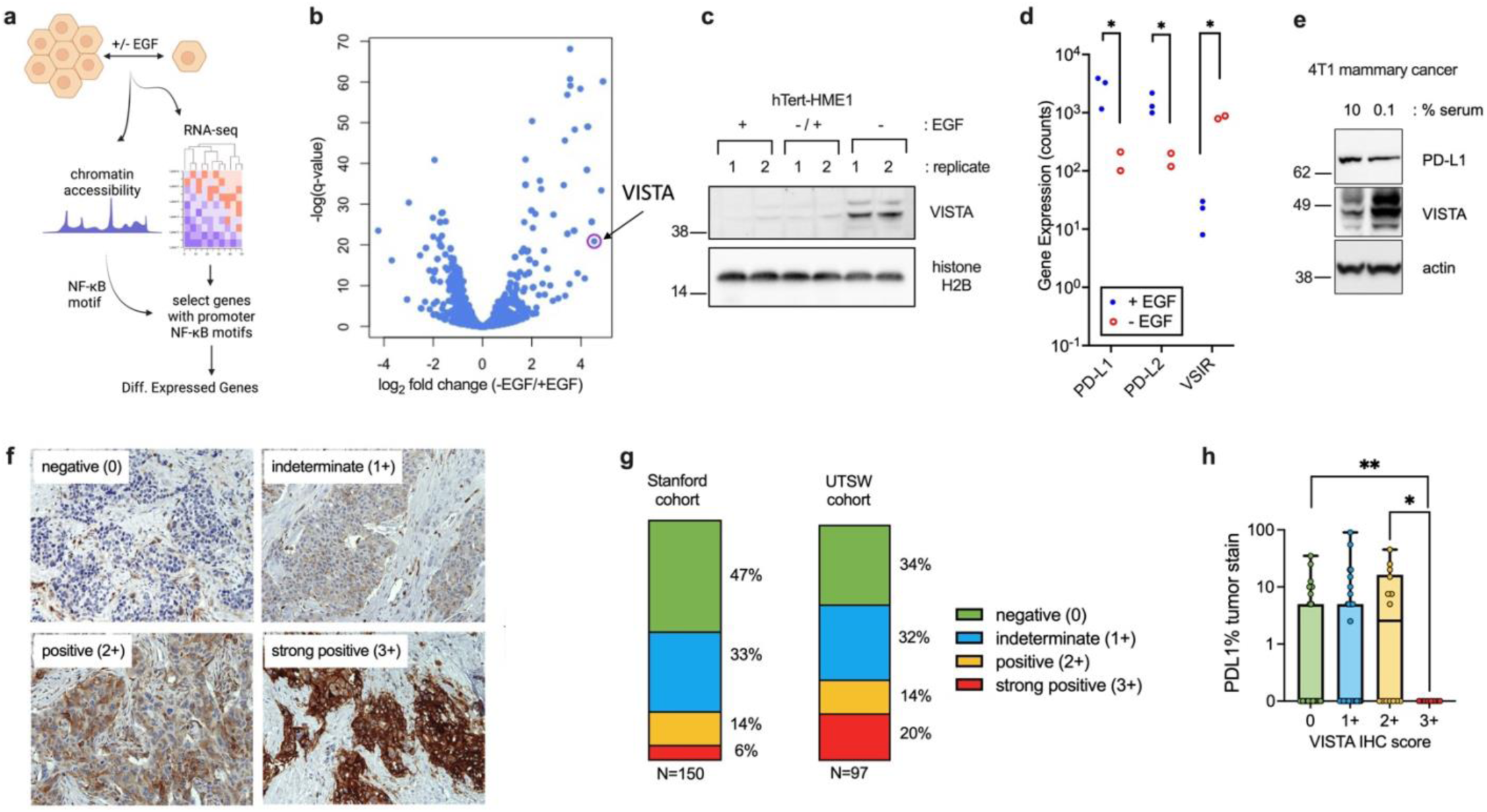
VISTA expression in mammary cell lines and human triple-negative breast cancers. **a.** Experimental and computational workflow to identify immunological target genes regulated in mammary cells. **b.** Chromatin accessibility and gene expression profiling were used to define transcripts with upstream NF-κB transcription factor binding motifs that were differentially expressed in the presence or absence of EGF. VISTA transcript is circled. **c.** Immunoblots of hTert-HME1 cells grown +EGF for 3 days, -EGF for 3 days, or -EGF for 2 days, then +EGF for 1 day (-/+ EGF). **d.** RNA-seq transcript levels for members of the B7 family in mammary cells grown +/- EGF. * q < 0.001, N= 3 replicates for +EGF and 2 replicates for -EGF. **e.** 4T1 cells were grown in 10% or 0.1% serum for 48 hours then protein levels were measured by immunoblot. **f.** Human TNBCs stained for VISTA and classified by VISTA score (0-3+) with higher value indicating stronger staining intensity of tumor cells. Magnification is 200x. **g.** Percentage of each VISTA score quantified in Stanford and UTSW cohorts (N=150 and 97, respectively). **h.** Quantification of tumor PD-L1 staining in Stanford TNBC cohort stratified by VISTA score. ** p < 0.01, * p < 0.03 by Welch’s t-test, or Welch ANOVA with Dunnett’s T3 multiple comparison test. n=103, n.s. = not significant

Because VISTA expression was detected in mammary cells, we next assessed a panel of 150 human primary TNBCs for VISTA expression by immunohistochemistry (IHC). Tumor cell staining for VISTA was detected in 20% of these tumors, of which 6% had extremely high levels with membrane staining (VISTA score = 3+; **Fig. 1f, g**). This was confirmed in an additional cohort of 97 tumors from a distinct patient population, in which 34% of tumors were VISTA positive, and 20% were strong positive (3+) (**Fig. 1g**). Tumors with the highest VISTA levels (3+) showed decreased tumor PD-L1 staining (**Fig. 1h**). Therefore, a subset of human TNBCs overexpresses VISTA and those with the highest VISTA levels lack PD-L1 expression.

The consequences of VISTA overexpression in TNBCs were modeled in human TNBC cell lines by expressing VISTA in cells with low basal VISTA levels. In HCC1806 TNBC cells the VISTA+ cells proliferated at a slower rate than WT parental or GFP control cells (**Fig. 2a**). Similarly, VISTA-expressing HCC1806 tumors showed significantly slower growth kinetics than controls (**Fig. 2b**). To confirm these results 4T1 mammary triple-negative cell lines were engineered to express VISTA, which also caused a significant growth defect in vitro compared to parental WT cells (**Fig. 2c**). When grown *in vivo* as orthotopic tumors in syngeneic mice VISTA+ 4T1 also grew at a slower rate compared to parental cells (**Fig. 2d**). Similar findings were observed in engineered VISTA+ EO771 tumors, compared to parental tumors, which had low VISTA expression (**Fig. 2e**). Therefore, in both immunodeficient and immunocompetent hosts, VISTA+ cancer cells had slower tumor growth, suggesting that this effect does not require a functioning immune system.

**Figure 2:**
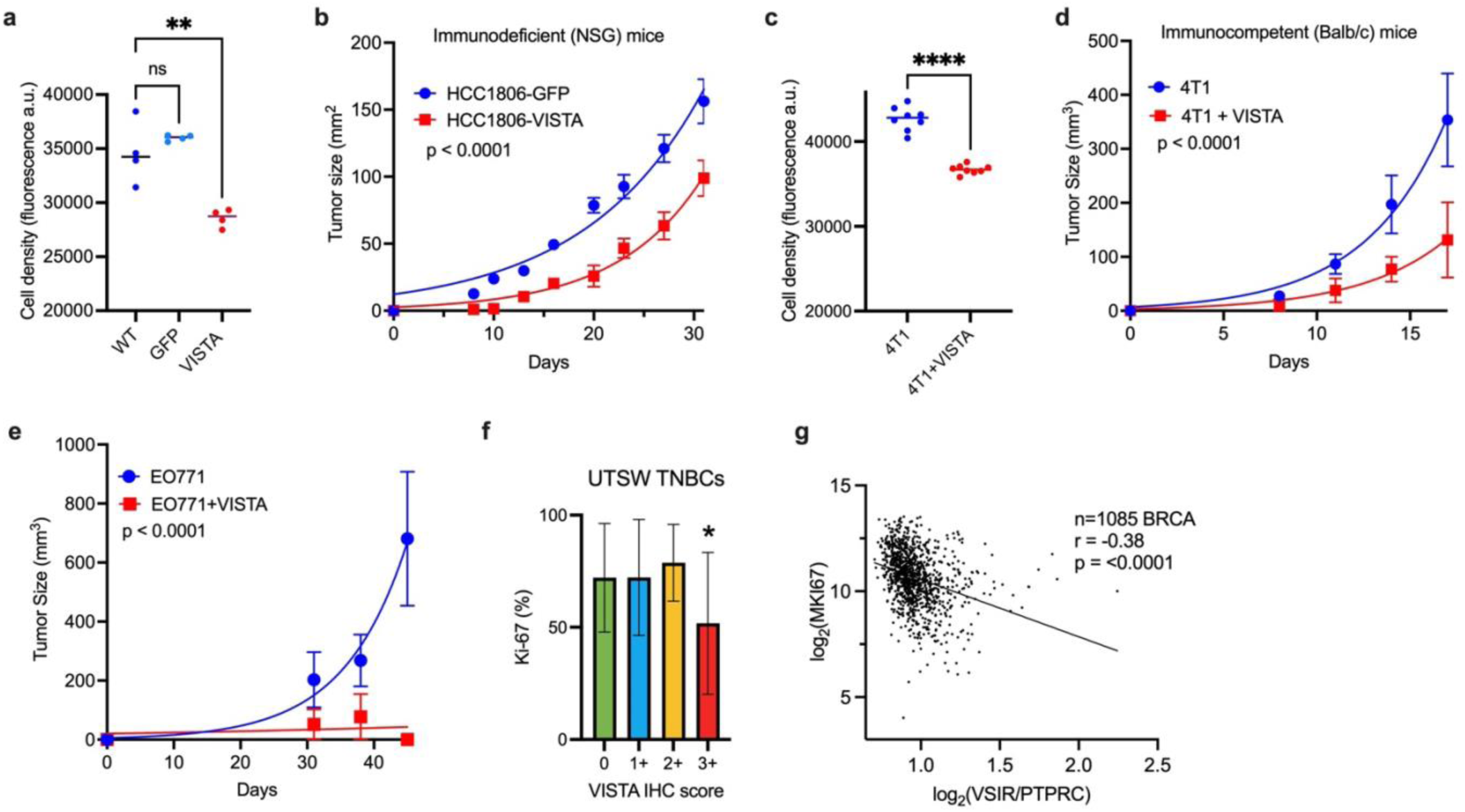
VISTA expression in TNBC causes diminished cell-intrinsic tumor growth. **a.** HCC1806 cells expressing GFP or VISTA were grown in serum and cell density was assessed by cell titer blue. ** p< 0.01 by ANOVA with Dunnett’s multiple comparison test (n=4 per group). **b.** GFP or VISTA-expressing HCC1806 cells were injected into mammary fat pads of immunodeficient NSG mice and tumor growth was measured. p-value by sum-of-squares F-test, n= 5 mice per group. **c.** 4T1 parental or VISTA+ cells were grown in culture and cell density was measured by cell titer blue. **** p< 0.001 by Welch’s t test (n=8 per group). **d.** Tumor growth curves of Balb/c mice bearing wt or VISTA+ 4T1 tumors. p-value by sum-of-squares F-test. n= 5 mice per group **e.** EO771 cells (WT or VISTA+) were grown in C57Bl/6 mice and tumor sizes were measured. p-value by F test. **f.** TNBC tumors from UTSW cohort (n=97) assessed for VISTA levels and Ki-67 score (%). * p = 0.011 by ANOVA. **g.** Breast tumors from TCGA (n=1085) were examined for gene expression of VSIR (normalized to PTPRC to adjust for presence of TILs) compared to MKI67. Correlation and p-value by Pearson’s. Correlation plotted as a solid line.

To extend these findings, human TNBC specimens were analyzed for Ki-67, a marker of active cell division (**Fig. 2f**). Whereas most TNBCs had high proliferative indices with Ki-67 scores of ∼75%, the tumors with the highest VISTA protein levels (3+ score) had significantly lower Ki-67 levels (∼50%). A similar relationship between VISTA and Ki-67 was observed in The Cancer Genome Atlas (TCGA) breast cancers (**Fig. 2g**). These findings confirm that breast cancer tumor cells with high VISTA levels have an intrinsically slower proliferative index.

### High VISTA expression alters EGFR receptor activation

Next, we sought to determine how VISTA affects cell-intrinsic proliferation. First, we characterized the correlation between VISTA levels in cell lines and tumor VISTA levels. In the TCGA BRCA dataset VISTA levels varied over 100-fold (**Fig. S1a**). Quantitative immunoblotting of VISTA+ versus WT HCC1806 cells showed that enforced expression of VISTA caused ∼100-fold increase in VISTA levels compared with starved WT cells (**Fig. S1b**). Therefore, enforced expression of VISTA mirrors VISTA levels in the highest TCGA tumors, whereas WT HCC806 cells expressed very low VISTA levels at baseline. VISTA expression also caused increased plasma membrane localization of VISTA (**Fig. 3a**), which was also evident in the VISTA 3+ tumors (**Fig. 1f**). Thus, VISTA+ HCC1806 cells accurately modeled features of VISTA-high tumors.

**Figure 3:**
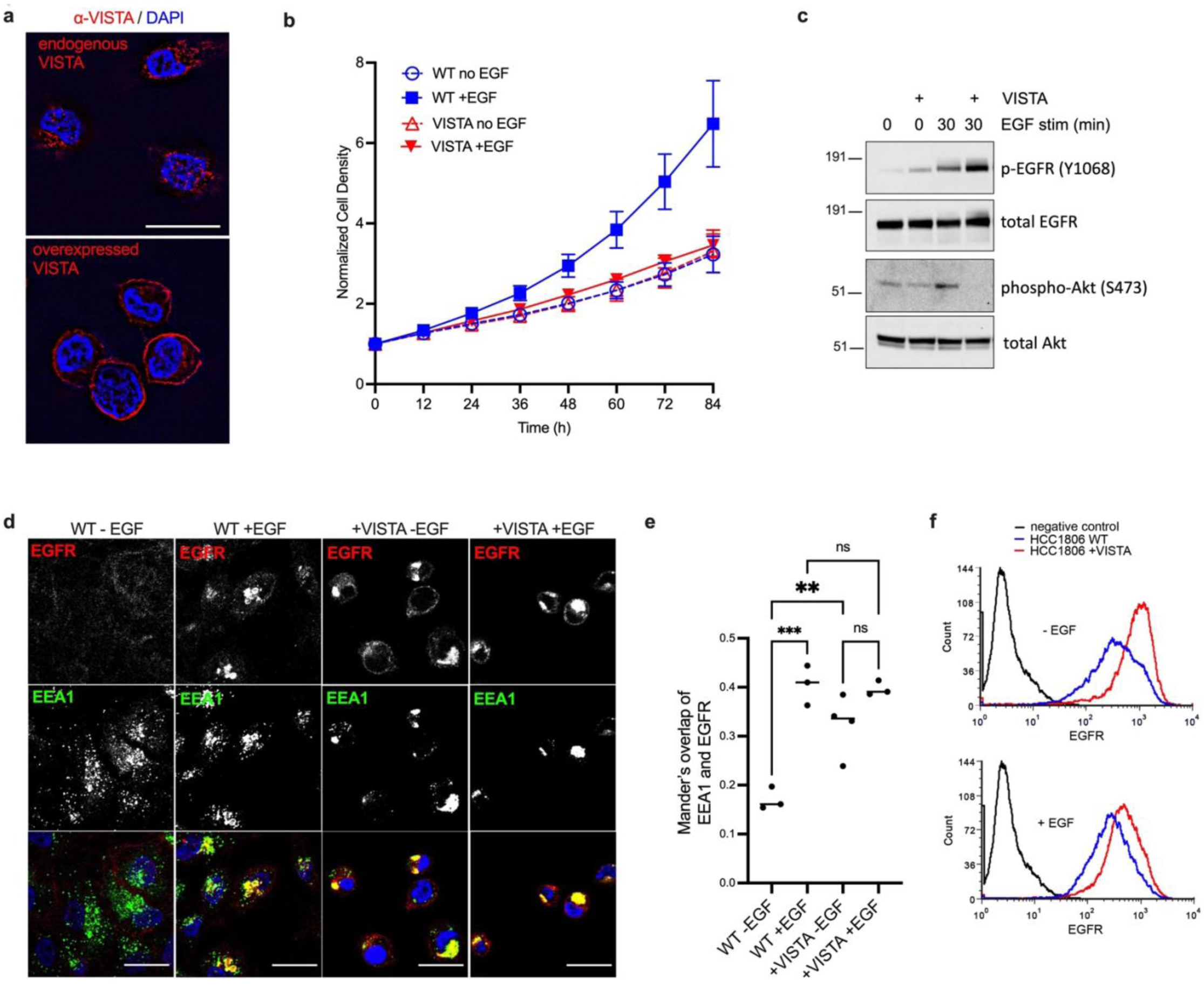
Enforced VISTA expression alters EGFR trafficking. **a.** Immunofluorescence detection of VISTA in WT (top-long exposure) and VISTA+ (below-short exposure) HCC1806 cells after serum starvation for 48 hours. Scale bar 20 μm. **b.** HCC1806 cells (WT or VISTA) were serum starved or stimulated with EGF and cell density was measured by Incucyte. **c.** HCC1806 cells were serum starved then stimulated with EGF for 30 minutes and proteins were analyzed by SDS-PAGE and immunoblot. **d.** Control (WT) or VISTA+ HCC1806 cells were serum starved for 48 hours then stimulated with EGF for 30 minutes. EGFR and EEA1 were detected by confocal microscopy. Scale bar 20 μm. All images are taken with the same exposure. **e.** Quantification of Manders’ overlap between EGFR and EEA1 from panel b (n=3 images per group, p< 0.001 by ANOVA). **f.** Flow cytometry of EGFR surface staining in unstimulated and EGF-stimulated HCC1806 cells.

Since VISTA+ tumors grew slow, we interrogated whether VISTA expression interfered with the response to growth factor stimulation. In cell proliferation assays, high VISTA levels caused diminished EGF-dependent growth compared to WT cells with low VISTA levels (**Fig. 3b**). This led us to test whether VISTA could disrupt EGFR signal transduction. Immunoblots for EGFR tyrosine 1068 phosphorylation, a site associated with receptor activation, demonstrated that VISTA+ cells had increased basal EGFR phosphorylation during starvation conditions compared to control cells (**Fig. 3c**). After EGF stimulation, VISTA+ cells continued to have increased EGFR phosphorylation, but this signal was not properly transmitted because Akt phosphorylation, which is critical for cell proliferation, was decreased compared to control cells (**Fig. 3c**). Therefore, EGFR signal transduction was decoupled by high VISTA levels.

De-phosphorylation of EGFR requires trafficking through endosomal compartments for exposure to phosphatases PTPRG/J and PTPN2, which determine sensitivity to growth factor and duration of signaling [36]. Therefore, we tested VISTA+ cells for alterations in EGFR trafficking. Control and VISTA+ cells were treated with serum starvation followed by EGF stimulation and then EGFR localization was assessed by confocal microscopy and flow cytometry. In serum starved control cells, EGFR had endosomal and plasma membrane localizations, as characterized by co-localization with EEA1 (an early endosome marker) and surface staining by flow cytometry. When stimulated by EGF control cells had increased EGFR co-localization with EEA1+ endosomes (**Fig. 3d, e**). However, in VISTA+ cells EGFR instead strongly co-localized with EEA1+ endosomes equally in unstimulated and EGF-stimulated cells (**Fig. 3d, e**). Also, VISTA+ cells had increased EGFR surface plasma membrane localization as detected by flow cytometry in both unstimulated and EGF-stimulated conditions compared to WT cells (**Fig. 3f**). Thus, VISTA expression altered EGFR localization by increasing EGFR residence at the plasma membrane and in EEA1+ endosomes.

The observed defect in EGFR trafficking suggested that other membrane trafficking processes could be altered in VISTA+ cells. However, VISTA+ cells did not have decreased macropinocytosis [37] (**Fig. S1c, d**) or transferrin receptor endocytosis compared to control cells (**Fig. S1e**), and instead both processes were slightly elevated in VISTA+ cells. Thus, despite a defect in EGFR trafficking, other membrane regulated processes remained robust in VISTA+ cells. Also, we tested whether EGFR was specifically affected by VISTA or whether other growth factor receptors were also altered. Exposure of cells to serum (which does not contain sufficient EGF to activate EGFR) consistently yielded decreased cell density of VISTA+ cells compared to parental cells (**Fig. S1f**, p<0.001 by F-test for WT vs. +VISTA, in both -/+ FBS conditions). Similarly, serum-induced phosphorylation of Akt was also decreased in VISTA+ cells compared to control, despite increased basal EGFR phosphorylation in VISTA+ cells (**Fig. S1g**). This indicates that VISTA may modulate other serum-responsive growth factor receptors, in addition to its role in modulating EGFR.

### Discovery of signaling adaptor proteins associated with VISTA CTD

The modulation of EGFR localization and activity by VISTA prompted us to determine the molecular basis of this effect. We focused on the intracellular C-terminal domain (CTD) of VISTA which lacks any defined modules or motifs. First, we expressed a mutant VISTA lacking the entire CTD (VISTA-ΔC; **Fig. S2a**). Upon EGF stimulation, VISTA-ΔC cells showed increased EGFR endocytosis and dissipation of plasma membrane-localized EGFR compared to cells expressing full-length VISTA (**Fig. S2b, c**). Next, mutants were constructed to sequentially delete 20 amino acid blocks through the CTD (**Fig. S2d**). All mutants expressed well (**Fig. S2e**), except for deletion mutant #4, which was not further studied. All VISTA mutants had lost some ability to restrict EGFR localization, in comparison to full-length VISTA (**Fig. S2f, g**). This suggests that multiple regions within the VISTA CTD may affect EGFR localization.

To identify potential VISTA CTD binding partners, the C-terminus of full-length VISTA was fused to the biotin-ligase TurboID (**Fig. 4a**). As a control, the VISTA-ΔC protein was fused to Turbo ID. These fusion proteins were expressed in HCC1806 cells, and the resulting biotinylated VISTA-associated proteins were recovered by streptavidin pulldown followed by mass spectrometry for peptide identification. Four proteins were specifically associated with the VISTA CTD: Rab11FIP5, Rab11FIP1, NUMB and GULP1 (**Fig. 4b**). These protein interactions were confirmed by repeating the streptavidin pull-down and immunoblotting with specific antibodies (**Fig. 4c**). Also, α-VISTA antibodies that bind to the extracellular domain were used to immunoprecipitate complexes. GULP1, NUMB and Rab11FIP1 were specifically enriched by VISTA antibodies in VISTA+ but not control cells (**Fig. 4d**). Thus, proximity biotinylation identified candidate proteins that interacted with VISTA-CTD.

**Figure 4:**
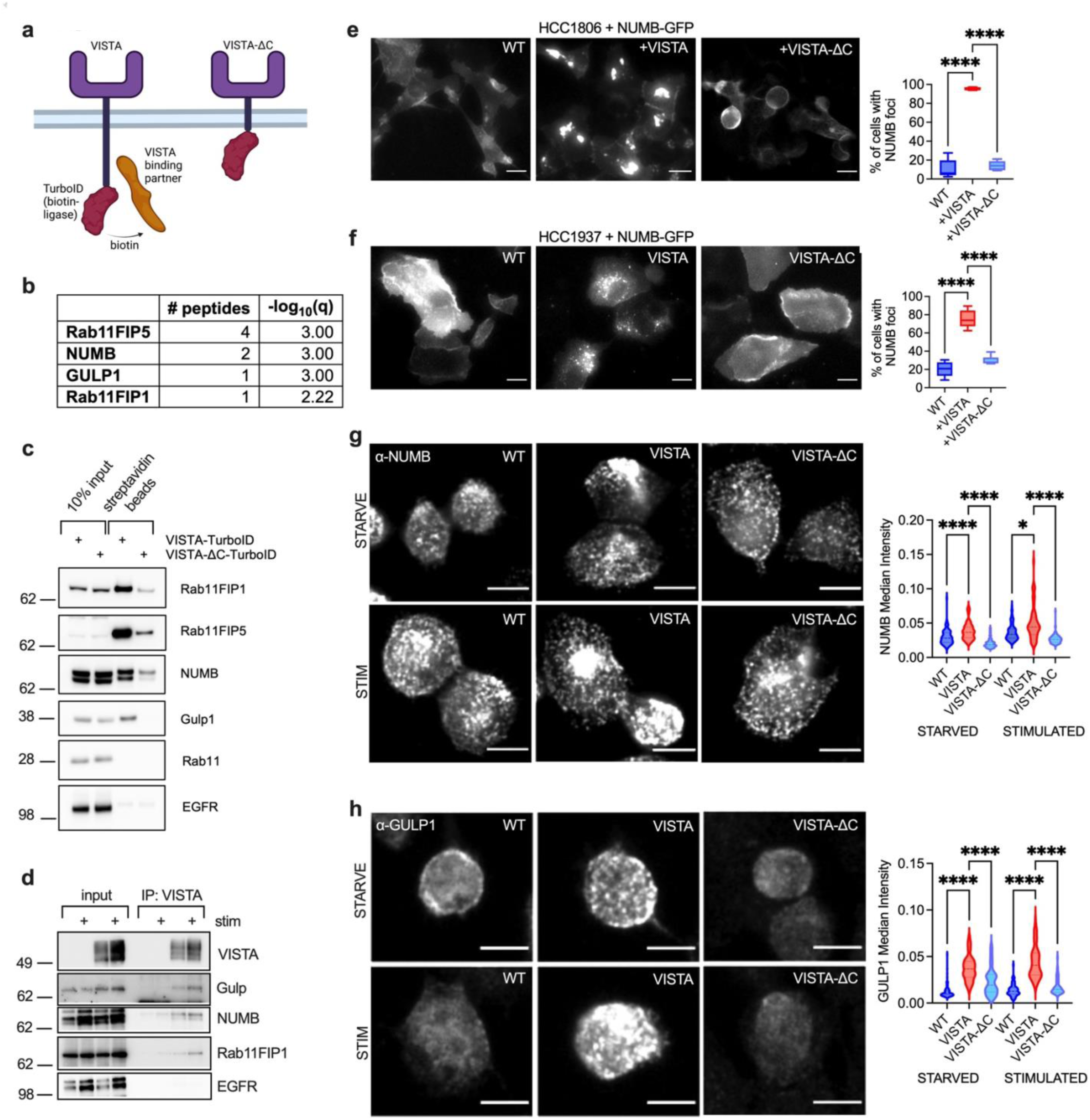
Identification of Rab11-FIPs and clathrin adapter proteins recruited to the VISTA CTD. **a.** Schematic of TurboID experiments with VISTA or VISTA-ΔC fusion proteins. **b.** Mass spectrometry of protein IDs of streptavidin-enriched unique peptides identified in the full-length VISTA-TurboID cells, but not in the VISTA-ΔC-TurboID cells. **c.** Streptavidin pulldowns followed by immunoblotting from VISTA-associated proteins in VISTA-TurboID and VISTA-ΔC-TurboID cells. **d.** WT or VISTA+ HCC1806 cells were serum starved for 48 h, or then stimulated with serum + EGF, then crosslinked with 0.2% formaldehyde and immunoprecipitated with α-VISTA antibodies, then detected by immunoblot. **e.** NUMB-GFP fusion protein was expressed in HCC1806 WT, VISTA, or VISTA-ΔC cells and imaged by fluorescence microscopy. Scale bar is 20 μm. (right) Percent of cells with NUMB-GFP foci within a field-of-view is plotted. N = 5 fields of view for each condition. **** q < 0.0001 by ANOVA with Tukey’s multiple comparisons test. **f.** NUMB-GFP fusion protein WT expressed in HCC1937 WT, VISTA, or VISTA-ΔC cells and imaged by fluorescent microscopy. Scale bar is 20 μm. **** q < 0.0001 by ANOVA with Sidek’s correction, N = 5, 8, or 6 fields-of-view per condition, respectively. **g.** HCC1806 WT, VISTA+, or VISTA-ΔC cells were serum starved for 48 hours then exposed to serum + EGF for 30 minutes, followed by immunofluorescence staining of endogenous NUMB. Median vesicle intensity is quantitated, N= 99, 99, 103, 132, 62, 107 vesicles, respectively. Significance by Kruskal-Wallis test with Dunn’s correction. ****p<0.0001, *p<0.04. **h.** Similar to c. with immunofluorescence staining of GULP1. N = 156, 80, 59, 91, 83, 99 vesicles, respectively.

The candidate VISTA-interacting proteins identified fell into two classes. First, both Rab11FIP5 and Rab11FIP1 are bona fide binding partners of Rab11, which controls receptor recycling [38]. The second class of proteins includes NUMB and GULP1, which are adaptor proteins that mediate receptor signaling and activity. Therefore, VISTA could potentially control EGFR activation by binding or modulating adapter proteins, such as NUMB, GULP1 or Rab11FIPs.

To investigate whether VISTA could modulate NUMB function, a GFP-tagged NUMB protein was studied. In WT cells, NUMB showed diffuse localization that was consistent with plasma membrane localization (**Fig. 4e**). However, in VISTA+ cells NUMB-GFP was detected in brightly stained intracellular foci (**Fig. 4e**). Interestingly, VISTA-ΔC did not cause NUMB-GFP to localize at intracellular foci (**Fig. 4e**). These experiments were repeated in an independent TNBC cell line, HCC1937, with nearly identical results (**Fig. 4f**). Similar effects were observed when probing for endogenous NUMB or GULP1 localization (**Fig. 4g, h**). Therefore, VISTA redirected receptor adaptor proteins to an endosomal compartment, and this effect required its CTD.

### VISTA recruits NUMB to EEA1+ early endosomes

Next, experiments were performed to clarify how VISTA caused NUMB recruitment to endosomes. A panel of various endosomal markers including caveolin, EEA1, LAMP1, Rab7 and Rab11 were visualized in cells that co-expressed VISTA and NUMB-GFP (**Supp. Fig. S3a**). Striking overlap was observed between EEA1 and NUMB-GFP foci compared to relatively less overlap observed with Rab11 (**Fig. 5a**). Confocal microscopy showed that VISTA co-localized with both Rab11+ vesicles and EEA+ vesicles (**Fig. 5b, c**). Taken together, these data suggest that VISTA can trap NUMB on EEA1+ vesicles, and that VISTA is distributed to both early and recycling endosomes.

**Figure 5:**
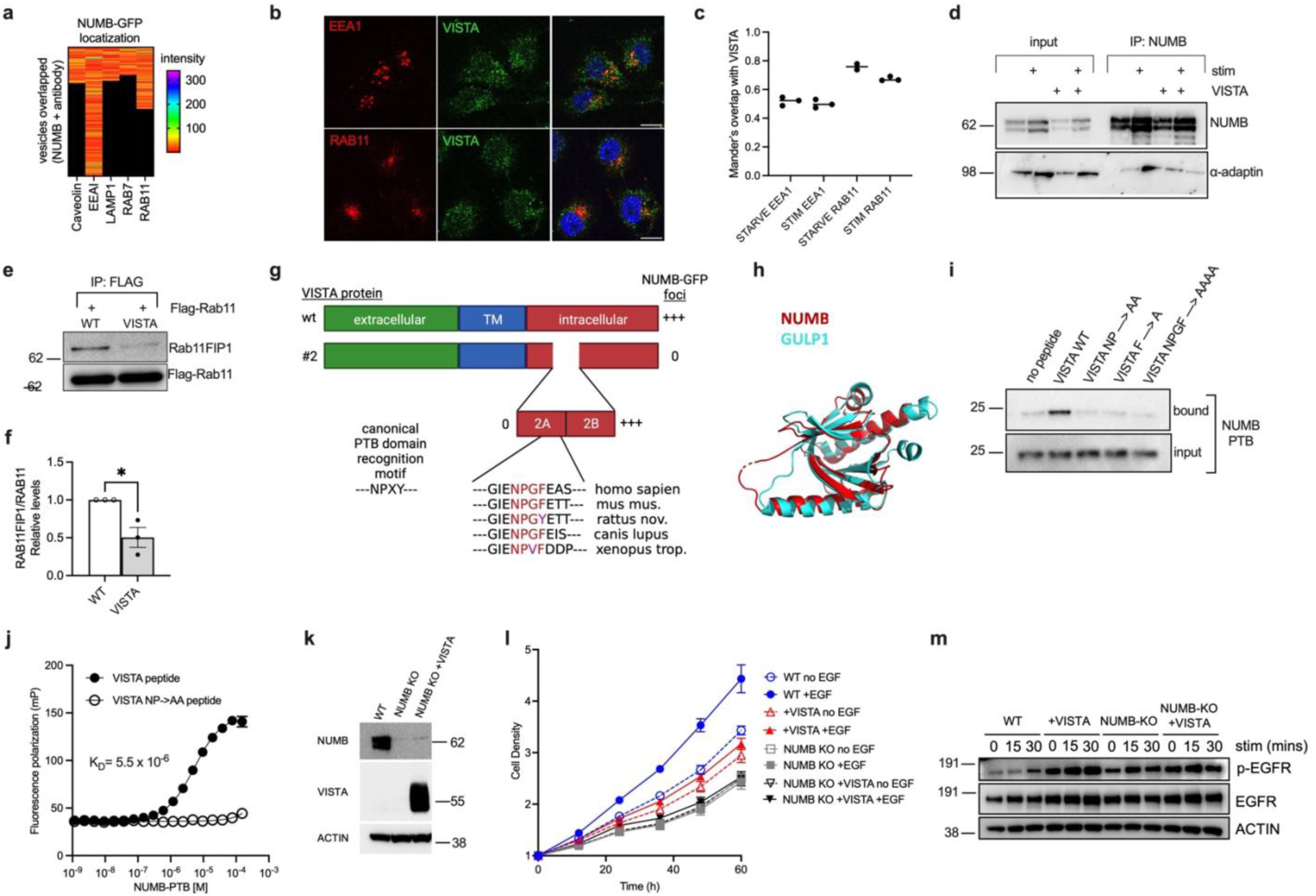
VISTA sequesters NUMB on EEA1+ vesicles. **a.** NUMB-GFP fusion protein was expressed in HCC1806 VISTA+ cells and co-localization was quantified for the indicated antibodies. Data represents all red signal within GFP+ vesicles from 3 independent photos at 63x resolution (N = 195, 154, 141, 176, 146 GFP+ vesicles for caveolin, EEA1, LAMP1, Rab7, Rab11, respectively). **b.** Confocal images of VISTA co-staining with EEA1 and Rab11. **c.** Co-localization of endogenous VISTA with Rab11 or EAA1 in serum starved and stimulated cells by Manders’ colocalization coefficient. N=3 pictures. **d.** Immunoprecipitation and immunoblotting of endogenous NUMB from HCC1806 cells +/- VISTA expression treated with serum starvation for 2 days, then stimulated (serum + EGF) for 30 minutes. **e.** WT and VISTA+ HCC1806 cells expressing Flag-Rab11 were subjected to Flag IP then immunoblotting. **f.** Quantitation of e. by densitometry. N=3 independent experiments. *p<0.02 by unpaired t-test. **g.** Schematic of VISTA protein sequence with sub-deletions in region 2 (2A, 2B) and sequence alignment of NPGF motif across various vertebrate species. **h.** 3D protein sequence alignment of crystalized PTB domains from NUMB (PDB: 5NJJ, red) and GULP1 (PDB: 6ITU, teal). **i.** VISTA 26-mer peptides centered on the NPGF motif were synthesized with and without mutations in NPGF and tested for interaction with NUMB-PTB proteins by streptavidin pulldown, followed by immunoblotting. **j.** Fluorescence polarization measurement of wt and mutant VISTA peptides binding to recombinant NUMB-PTB protein. K_D_ calculated for wt peptide binding to NUMB-PTB. **k.** Immunoblot of HCC1806 cell lines engineered with CRISPR knockout of NUMB and NUMBL, and VISTA expression. **l.** Proliferation measurement of HCC1806 cell lines by Incucyte in the presence or absence of EGF. **m.** Immunoblots of cells grown without serum or EGF for 2 days, then stimulated with serum and EGF for the indicated times.

Next, we tested whether NUMB sequestration at endosomes interfered with NUMB binding the AP2 subunit of clathrin, which typically occurs at the plasma membrane. In WT cells, NUMB co-immunoprecipitated with the AP2 subunit α-adaptin after growth factor stimulation (**Fig. 5d**). However, in VISTA+ cells, the association between NUMB and AP2 diminished (**Fig. 5d**). Therefore, VISTA sequestered NUMB at early endosomes, which likely prevents its association with regulators at the plasma membrane.

To further investigate how VISTA blocked NUMB trafficking in EEA1+ vesicles, we hypothesized that VISTA disrupted the association between Rab11 and Rab11FIP proteins, thereby preventing effective trafficking of NUMB. To test this, FLAG-Rab11 was expressed in WT and VISTA+ cells, and FLAG immunoprecipitation was performed. The amount of Rab11FIP recovered with Rab11 was significantly decreased in VISTA+ cells compared to WT cells (**Fig. 5e, f**). This indicated that VISTA may prevent association between Rab11 and Rab11FIP proteins, which could disrupt NUMB localization.

### Identification of a VISTA intracellular motif required for NUMB redistribution

To further define the mechanism by which VISTA could trap NUMB on early endosomes, we mapped regions of the VISTA CTD required to cause NUMB foci formation (**Fig. S3b, c**). Only deletion 2 completely blocked NUMB foci formation (**Fig. S3b**). Region 2 contained a conserved NPGF sequence (**Fig. 5g**), which was similar to an N-P-X-pY motif (X = any amino acid, pY = phosphotyrosine) that is known to bind phosphotyrosine-binding (PTB) domains and does not strictly require pY [39]. Both VISTA binding partners NUMB and GULP1 contain PTB domains with extensive 3D overlap (**Fig. 5h**). Thus, the NPGF motif in VISTA was a good candidate for a NUMB-binding region via its PTB domain.

To evaluate whether the VISTA NPGF motif could bind NUMB, we synthesized a 26-residue biotinylated peptide centered on the NPGF motif. In streptavidin pulldown assays, this peptide enriched recombinant NUMB PTB domain (**Fig. 5i**). Next, a series of peptides was synthesized with alanine mutations to one, two or all four NPGF residues. All mutations in the NPGF motif disrupted NUMB binding (**Fig. 5i**). This was further confirmed in a fluorescence polarization assay that showed specific binding between NUMB-PTB to VISTA WT peptide, but no binding to a mutated peptide (**Fig. 5j**). This assay allowed us to calculate a K_D_ of 5.5 μM (95% CI 5.1-6.0 μM) for VISTA NPGF binding to the NUMB-PTB. Taken together, these studies provide strong evidence that the VISTA NPGF motif binds directly to NUMB.

Next, we sought to confirm that NUMB binding to VISTA played a functional role in EGFR activation in breast cancer cells, as suggested by prior studies in neural systems [40]. We generated an HCC1086 cell line with targeted CRISPR deletions of both NUMB isoforms (NUMB and NUMBL) and expressed VISTA in the background of NUMB knockout (KO; **Fig. 5k**). These cell lines were used for proliferation assays in the presence or absence of EGF stimulation. NUMB KO cell lines had diminished cell proliferation and were not sensitive to EGF stimulation, compared to parental cells (**Fig. 5l**). Expression of VISTA in the NUMB KO background did not further impair proliferation. Thus, NUMB was required for EGF-induced proliferation and the anti-proliferative effect of VISTA required NUMB.

Next, NUMB KO cells were examined for EGFR localization and trafficking to further clarify the role of NUMB in cell proliferation. NUMB KO and WT cells showed similar levels of EGFR/EEA1 co-localization (**Fig. S3d, e**). This indicated that NUMB is not required for EGFR trafficking to EEA1+ endosomes. However, NUMB KO cells had increased basal EGFR phosphorylation compared to WT cells, which was similar to increases in EGFR phosphorylation observed in VISTA+ cells (**Fig. 5m**). This suggests that NUMB primarily functions to resolve EGFR phosphorylation, likely by directing EGFR through downstream endosomal compartments, which is known to be required for proper EGF-induced signal transduction [36].

### The NPGF motif is required for VISTA function

We further dissected the function of NUMB binding to VISTA by constructing two VISTA mutants in which the NPGF residues were either deleted or mutated to four alanine residues. VISTA expression and localization were then examined by confocal microscopy and flow cytometry. Both techniques demonstrated stronger plasma membrane expression of WT VISTA, compared to VISTA NPGF deletion or mutant proteins (**Fig. 6a, b**). Intracellular flow cytometry showed that VISTA mutants were retained to a higher degree in an intracellular compartment than WT VISTA (**Fig. 6c**). A third mutant in which the intracellular domain of VISTA was swapped with the PD-L1 intracellular domain (which lacks a NUMB-binding motif) mirrored the results of NPGF mutation (**Fig. 6b, c**). Thus, the NPGF motif affected VISTA localization to membrane compartments.

**Figure 6:**
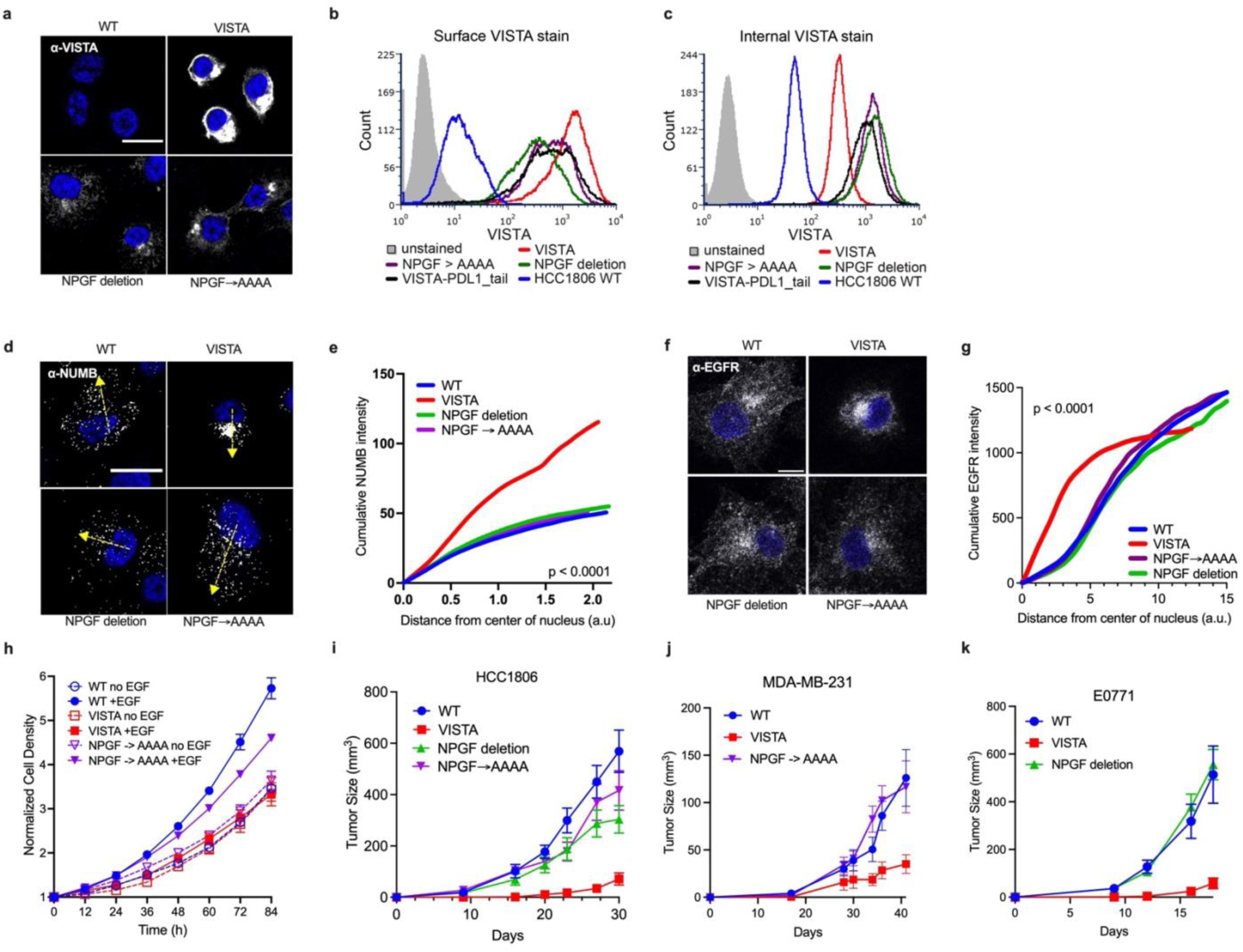
An NPGF domain in VISTA CTD controls NUMB localization and tumor growth. **a.** Confocal microscopy images of VISTA localization in HCC1806 cells expressing the indicated VISTA proteins. Scale bar = 20 μm, DAPI is blue. **b.** Surface flow cytometry with an antibody towards VISTA of HCC1806 cells expressing the indicated VISTA proteins. **c.** HCC1806 cells were subjected to trypsinization, fixation, permeabilization and flow cytometry with antibodies towards VISTA. **d.** Confocal microscopy of endogenous NUMB localization in HCC1806 cells expressing VISTA proteins. Scale bar = 25 μm. Yellow arrows indicate vector of intensity quantification. **e.** Quantification of NUMB intensity along a vector from the center of the nucleus to the cell periphery (yellow arrows). p value by F test, n=10 cells per conditions. **f.** Confocal microscopy of endogenous EGFR localization in HCC1806 cells expressing VISTA proteins. Scale bar = 20 μm. **g.** Quantification of EGFR intensity along a vector from the center of the nucleus to the cell periphery. p value by F test, n=10 cells each. **h.** Incucyte cell density data of HCC1806 cells grown in serum-free media with or without EGF addition (n=3 wells). **i.** Tumor growth measurements of NSG mice injected in mammary fat pads with HCC1806 cells expressing VISTA proteins, n=5 mice (10 tumors) per cohort. **j.** Tumor growth measurements of NSG mice injected in mammary fat pads with MDA-MB-231 cells expressing VISTA proteins, n=5 mice (10 tumors) per cohort. **k.** Tumor growth measurements of C57Bl/6 mice injected in mammary fat pads with E0771 cells expressing VISTA proteins, n=5 mice (10 tumors) per cohort.

Next, the functional impact of VISTA NPGF mutations on NUMB and EGFR localization was assessed. Unlike full-length VISTA, the NPGF mutants did not alter NUMB or EGFR distribution throughout the cytoplasm (**Fig. 6d-g**). In proliferation assays, mutation of NPGF to AAAA restored EGF sensitivity, in contrast to full-length VISTA, which rendered cells insensitive to EGF (**Fig. 6h**). Next, these VISTA mutants were assessed in three orthotopic models of triple-negative breast cancer including two immunodeficient xenografts (HCC1806 and MDA-MB-231) and one syngeneic transplantable model (EO771). In all models either alanine mutations or deletions of the NPGF motif restored tumor growth to near WT levels, whereas expression of full-length VISTA impaired tumor growth (**Fig. 6i-k**). Taken together, VISTA’s NPGF motif was essential for growth and protein localization phenotypes conferred by VISTA in tumor cell lines.

### Physiologic VISTA levels control EGFR and NUMB in breast cancer cells

Next, we assessed whether physiologic VISTA levels affected NUMB and EGFR localization by generating VISTA knockout (KO) triple-negative breast cancer cell lines (**Fig. 7a**). This revealed that VISTA KO cells had consistently decreased co-localization of EGFR with EEA1 compared to control or VISTA-mutant cells after EGF stimulation (**Fig. 7b, c**). This effect was also observed in hTert-HME cells treated with shRNAs to deplete VISTA levels (**Fig. 7d-f**). Thus, physiological VISTA levels were required to promote co-localization of EGFR and EEA1 after growth factor stimulation.

**Figure 7:**
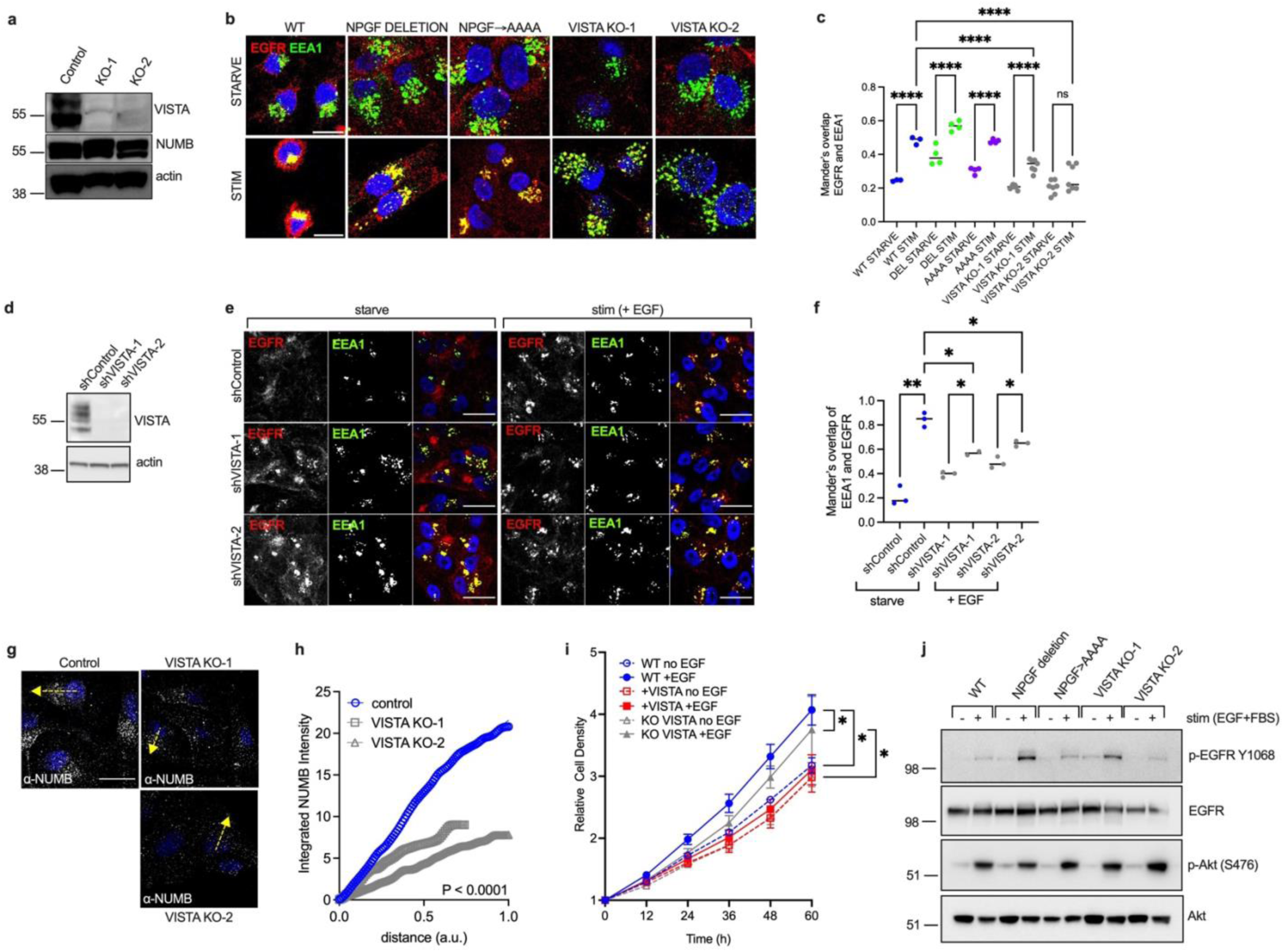
Physiologic VISTA levels control EGFR and NUMB in breast cancer cells. **a.** Immunoblots of protein levels in control and VISTA knockout (KO) HCC1806 cell lines. **b.** Confocal microscopy images of NPGF-mutant and VISTA KO cell lines starved or stimulated with EGF for 30 minutes. Scale bar = 20 μm. **c.** Quantification of images in b. EGFR and EEA1 overlap by Manders’. Each data point is from a single image (n=3-7 images). **** p<0.001 by ANOVA. **d.** Immunoblots of VISTA levels after shRNA treatments. **e.** Confocal images of EGFR and EEA1 localization in hTert-HME1 cells starved or stimulated with EGF for 30 min. Scale bar =30 μm. **f.** Quantification of images in e. EGFR and EEA1 overlap by Manders’. Each data point is a single image. * p < 0.05 by ANOVA. **g.** Confocal microscopy images of NUMB in HCC1806 cells with VISTA KO. Yellow line indicates vector of signal quantification. Scale bar = 30 μm. **h.** Quantification of integrated NUMB signal intensity along a vector from center of nucleus to edge of cell. n=10 cells per group. p value by F-test. **i.** Cell proliferation +/- EGF stimulation of HCC1806 cells (WT, VISTA+, VISTA KO) performed by Incucyte. *p < 0.01 by F-test. **j.** SDS-PAGE and immunoblot analysis of HCC1806 cells expressing mutant forms of VISTA, or with VISTA KO. Cells were serum starved for 2 days, then stimulated with serum + EGF for 30 minutes.

To determine how VISTA promoted EGFR localization to early endosomes, we examined NUMB localization in VISTA KO cells. Compared to control parental cells, VISTA KO cells had more dispersed NUMB localization under starvation conditions (**Fig. 7g, h**). In addition, although VISTA KO cells retained sensitivity to EGF stimulation, they were not as sensitive to EGF as parental cells, and EGFR phosphorylation was preserved in VISTA KO cells (**Fig. 7i, j**). This suggests that physiologic levels of VISTA promote NUMB to endosomes to promote EGFR trafficking, but that higher VISTA levels may be required to interfere with EGFR signaling.

## Discussion

Mechanisms of VISTA function remain obscure, despite a large body of data suggesting that targeting this receptor could be beneficial for cancer immunotherapy. In a search for novel immune targets for triple-negative breast cancer, we observed that a fraction of these tumors had high VISTA levels and diminished proliferative index compared to tumors with lower VISTA levels. Modeling in breast cancer cell lines and tumors confirmed that anti-proliferative effects of VISTA required its intracellular domain, and we identified the first functional intracellular motif (NPGF) required for these anti-proliferative effects. In breast cancer cell lines growth suppression is likely independent of the previously identified VISTA ligands PSGL-1 and VSIG3 because a functional immune system was not required for tumor suppression by VISTA. Instead, the data point to a role for VISTA in coordinating cell-intrinsic intracellular signaling of growth factor receptors, which include EGFR in tumor cells, but possibly other receptors as well.

VISTA appears to exert its effect on other receptors through its NPGF motif that recruits the adaptor protein NUMB, and likely other PTB domain-containing proteins, like GULP1. Previous studies have shown that NUMB helps to terminate receptor signaling via post-endocytic sorting or recycling [40–43]. This agrees with our findings that VISTA recruits Rab11FIP proteins. Also, we show that high VISTA levels disrupt Rab11 binding to Rab11FIP1, which should impair receptor recycling from endosomes back to the plasma membrane. Concordantly, VISTA is primarily localized to endosomes in breast cancer cells, whereas surface expression of VISTA is quite low under physiologic conditions in these cells. Taken together, these findings suggest that VISTA binds and sequesters NUMB at endosomes to repress proper receptor activation. Mutation of the NPGF motif is sufficient to release NUMB, which then binds growth receptors to promote signaling. Thus, by titrating NUMB location and activity, VISTA can control the function of various other receptors.

A large body of evidence points to positive roles for NUMB in receptor activation through various mechanisms, including recruitment of receptors to clathrin-coated pits [44, 45]. Although EGFR endocytosis was blocked by VISTA in breast cancer cells, this was not due to NUMB because NUMB knockout cells had no such endocytic defect, despite having increased signaling. Instead, the data argue for a more direct role for NUMB in controlling receptor activation at early stages. It is also possible that proper receptor recycling through endosomal compartments controlled by NUMB is required to prime receptors for re-activation. Nevertheless, the data suggest that a major molecular mechanism of VISTA effector function is to repress NUMB.

Deciphering molecular mechanisms allows for the design of therapeutic strategies. Common mechanisms of immune checkpoint control include binding or titration of co-receptors important for immune cell activation, or propagating negative signals through well-defined intracellular domains, like immunoreceptor tyrosine-based inhibitory motifs (ITIMs) [46, 47]. Although VISTA lacks conventional intracellular signaling domains, the NPGF domain identified here and the ability of VISTA to influence activation of cell surface receptors implies that VISTA binding to NUMB is critical for its ability to modulate signaling and cell activation status. This defines a previously unrecognized mechanism of receptor repression that could apply to immune checkpoint control and inform new therapeutic strategies. For example, to date all VISTA-targeting small molecules and antibodies block extracellular binding sites, which can lead to agonistic or antagonistic effects [16, 48–56]. It is possible that cell-penetrating small molecule strategies designed to target the intracellular domain of VISTA, for example by binding the NPGF motif to disrupt recruitment of NUMB or Gulp1, could be developed for anti-tumor immune checkpoint blockade for VISTA-negative tumor cells. Furthermore, our results predict that altered co-receptor activation or trafficking could be cellular outputs utilized to validate VISTA neutralization in pre-clinical or clinical studies. These insights should guide the design and application of future immune-targeting anti-cancer therapies.

## Acknowledgements

The authors would like to thank members of the Gruber, Snyder and Levy labs, especially Ron Levy, for helpful comments and suggestions during this work. We would also like thank to Marcel Mettlen for sharing reagents and protocols. This project was supported by a Susan G. Komen Postdoctoral Fellowship (PDF17483383) and an ASCO/Conquer Cancer Foundation Young Investigator Award sponsored by the Breast Cancer Research Foundation to J.J.G. J.J.G. was also supported by the Jane Coffin Childs Memorial Fund for Medical Research, the NIH (R01CA290297), CPRIT (RR200090) and the V Foundation (V2022-022). J.J.G. and M.L.T. acknowledge support from Stanford Cancer Institute. J.M.F. and M.P.S. acknowledge research support from NIH/NCI (U2CCA233311). Graphics were created with BioRender.com.

## Author Contributions

Y.Z., F.C., P.R., K.P., H.G., C.G., T.A., A.H., S.M., B.S.G., I.B-S., and J.J.G. performed experiments. J.J.G., T.Y., A.M.L., and M.P.S. designed the study and oversaw study implementation and data integrity. J.J.G., M.L.T., and M.P.S. obtained research funding. J.J.G., A.M.L., J.D.B.K, and D.T. wrote the manuscript. M.M.J., S.Y., C.L., V.E., A.W.K., Y.P., J.M.F., R.B.W performed specimen collection, staining or pathological analyses.

## Materials and Correspondence

Please direct correspondence and requests for resources and reagents to the Corresponding Authors Joshua Gruber and Michael Snyder.

## Data and code availability

All data that support the findings of this study are available in the manuscript and supporting information. There are no restrictions on data sharing. Custom code used to generate conclusions in this work will be available for download at https://github.com/GruberLabUTSW/. There are no restrictions on code availability.

## Methods

### Cell Lines

All cell lines were obtained from ATCC and cultured in a humidified incubator at 37°C, with 5% CO_2_. Cell lines E0771 (mouse), 4T1 (mouse), MDA-MB-231 (human female), HCC1937 (human female) and HCC1806 (human female) were maintained in RPMI with 10% FBS and 1% penicillin/streptomycin. hTert-HME1 was maintained in MEGM media (Lonza) with 1% penicillin/streptomycin. Cell lines were authenticated by morphology and STR analysis.

### Human tumor specimens

Informed consent was obtained prior to the acquisition of all clinical samples. Stanford Institutional Review Board and UTSW IRB approved all human research. Inclusion criteria required a diagnosis of triple-negative breast cancer (ER <10%, PR<10% and HER2 negative) for primary breast cancer specimens. Prior chemotherapy was allowed, but not required. All subjects were female and >18 years of age. Sample size was predetermined by existing tissue microarrays.

### Sex as a biological variable

For breast cancer studies, only female tumor specimens and female mice were used because the disease is only found in females. T cell experiments were performed in male and female animals with similar findings reported for both sexes.

### Lentiviral infections, immunofluorescence, cell assays, transferrin uptake

Lentivirus was produced in HEK-293T cells by co-transfection with pVSV-g and psPAX2 together with lentiviral gene expression vectors. Lentivirus was harvested and concentrated with Lenti-X reagent (Takara). Viral infections were performed with polybrene (8 μg/mL), then stable cell lines were selected with puromycin or blasticidin. **Immunofluorescence:** cells were plated on coverslips in 24-well plates, fixed in 4% formaldehyde for 15 minutes, permeabilized for 15 minutes with 0.2% triton-X-100 in PBS and blocked with 1.5% FBS. Primary antibody dilutions ranged from 1/100 to 1/200 and secondary Alexa Fluor antibodies were used at 1/200 dilution. Imaging was performed with a Keyence BX-800 microscope or a Leica fluorescence upright microscope or a Zeiss LSM 700 confocal microscope. **TMR-dextran uptake**: Cells were plated on coverslips, washed 3x with PBS, incubated in serum-free RPMI media overnight or media with 10% serum. Then, TMR-dextran at a concentration of 10 mg/ml was diluted to 1 mg/ml in growth media and added to wells for 30 minutes at 37°C, then washed 5x with PBS, fixed in 4% formaldehyde, mounted with DAPI, then imaged. Cell density was measured with CellTiterBlue, and fluorescence read on a Tecan Infinite M1000 plate reader. **Transferrin uptake**: Cells were seeded in 96 well plates for 24 h, then washed with PBS, then total and blank wells were fixed in 4% PFA for 10 min at RT and washed with PBS. Transferrin receptor antibody D65 (gift from Marcel Mettlen) was used at 1 μg/mL in serum-free media for the indicated timecourse at 37°C, then placed at 4°C to stop membrane trafficking, washed with cold acetic acid (11.483 mL glacial, 11.96 g NaCl in 1L ddH_2_O, pH 2.3) rapidly 4 times, then with cold PBS 4 times and fixed in cold 4% PFA for 1 minute then transferred to 37°C for 30 minutes. Subsequent processing included PBS washing, permeabilization in 0.1% Triton X-100 in PBS for 5 minutes at RT, PBS washing, blocking in 5% milk in PBS, PBS washing, incubation with 1:5000 dilution of anti-mouse-HRP conjugate overnight at 4°C, PBS washing, then development with Amplex Red, quenching with 5 M H_2_SO_4_ and detection at OD490 and OD650. Then cells were washed in PBS and cell density was detected by BCA assay for per-well normalization.

### Immunoblots, Microcapillary Assays, Flow Cytometry

Protein extracts were made in RIPA buffer and quantitated by BCA assay and diluted to equal concentrations. Polyacrylamide gel electrophoresis was performed on NuPAGE Novex gradient gels (Thermo Fisher) followed by wet transfer to nitrocellulose membranes. Blocking was briefly performed with 5% non-fat milk and primary antibody was incubated overnight at 4°C, followed by incubation with HRP-conjugated secondary antibody (Cell Signaling) at room temperature for 1 hour followed by washing, then developed with ECL pico or femto (Thermo Fisher).

Microcapillary immunoassays were performed with a ProteinSimple Wes machine. Surface flow cytometry was performed from adherent cells collected with EGTA/EDTA detachment to preserve surface antigens, followed by Fc-blocking with 1 μg of mouse IgG per 1 million cells (15 min incubation at RT in FACS buffer), followed by primary and secondary antibody staining and analysis on a FACS Calibur machine. For detection of intracellular antigens, cells were collected with trypsinization, fixed in 4% PFA for 15 minutes at RT, then permeabilized in 0.3% triton X-100 in FACS buffer for 10 minutes at RT, then proceeded to antibody staining and analysis as above.

### Streptavidin pulldowns

For streptavidin pulldown of biotinylated proteins, 8 x 10 cm plates of miniTurboID-expressing cells were grown in complete media supplemented with 500 μM biotin for one hour. Cells were lysed in 1 mL RIPA + protease inhibitors, sonicated, clarified by centrifugation and proteins were captured with 20 μL of pre-washed Streptavidin C1 MyOne Dynabeads (Thermo Fisher) for 3 hours at 4 °C. Then beads were washed five times in RIPA buffer, once in PBS, then resuspended in 0.1% SDS and 5 mM TCEP, then boiled for 3 minutes with lid caps. Samples were alkylated with iodoacetamide, then precipitated with ToPREP to remove interfering substances and resuspended in triethylammonium bicarbonate buffer. Trypsin was added and samples digested overnight, then desalted with C18 columns and dried in a vacuum centrifuge and processed for mass spectrometry. For peptide pulldowns of recombinant NUMB PTB domain peptides were synthesized with the n-terminal biotin-AHX linkages to the following sequences: (WT) RMDSNIQGIENPGFEASPPAQGIPEAK, (NP>AA) RMDSNIQGIEAAGFEASPPAQGIPEAK, (F>A) RMDSNIQGIENPGAEASPPAQGIPEAK, and (AAAA) RMDSNIQGIEAAAAEASPPAQGIPEAK. Recombinant NUMB was purified from BL21 DE3 e. coli transformed with NUMBLA plasmid (Addgene #42420, RRID Addgene_42420; a gift from Nicola Burgess-Brown) induced at O.D. 0.4-0.8 with 1 mM IPTG at 16°C overnight in 50 mL culture with chloramphenicol and kanamycin. Purification was done by sonication for 15 minutes on ice (15 sec on/30 sec off) with 60% amplitude on a Branson sonicator in the presence of lysis buffer (50 mM Tris, 300 mM NaCl), 14.2 mM β-ME and protease inhibitor cocktail tablet (Roche). Extract was centrifuged at 13,000 RPM for 30 min at 4°C, then supernatant was mixed with washed Ni-NTA resin (0.5 mL resin per 50 mL extract) for 1 h at 4°C, then transferred to drip column for washing (50 mM Tris, 300 mM NaCl, 25 mM imidazole, 5 mM β-ME), followed by elution (50 mM Tris, 300 mM NaCl, 300 mM imidazole, 5 mM β-ME, +/- 10% glycerol) and concentrated over 10K MCWO Amicon columns. Purified protein was dialyzed against dialysis buffer (50 mM Tris, 100 mM NaCl, 5 mM β-ME, 10% glycerol). Peptide pulldowns of recombinant NUMB PTB domain were performed in 1 mL JS Buffer (50 mM HEPES, 100 mM NaCl, 1% glycerol, 1% triton X-100, 1.5 mM MgCl_2,_ 5 mM EGTA), with streptavidin C1 magnetic beads (10 μL), 1 μL of (NUMB PTB 1 mg/mL), 1 μL of VISTA peptides (1 mg/mL) at 4°C for 1 hour, then washed with binding buffer and eluted in SDS-based gel loading buffer for 3 minutes at 100°C. Bound NUMB was detected by anti-His immunoblot after SDS-PAGE.

### Mass spectrometry

LC-MS/MS chromatography was performed by ITSI Biosciences (Johnstown, PA, USA) with a Thermo EASY-nLC system operating in the nano-range. Peptides were eluted from the column with a linear acetonitrile gradient from 5-32% over 90 minutes, followed by high and low organic washes for another 20 minutes into a Q-Exactive mass spectrometer (Thermo Fisher) via a nanospray source with the spray voltage set to 2.0 kV and the ion transfer capillary set at 250 °C. A data-dependent Top 15 method was used for a full MS scan from *m/z* 350-1600 followed by MS/MS scans of the 15 most abundant ions. Each ion was subjected to HCD (Higher energy C-trap dissociation) for fragmentation and peptide identification. Raw data files were searched against the most recent Human UniProt database using Proteome Discoverer 2.2 (Thermo Scientific) and the Sequest HT search algorithm. For protein identification and PTM results only peptides identified with high confidence were used. Tryptic fragments allowed for up to two missing cleavages per peptide. Oxidation of methionine and n-terminal acetylation were used as dynamic modifications, and carbamidomethyl of cysteine and biotin-lysine were used as static modifications. Target decoy was used for Peptide Spectrum Matches validation.

### Fluorescence polarization assay

Reactions were performed in Corning #3820 384 well assay plates in a CLARIOstar Plus instrument set for excitation at 482-16 nm, emission at 530-40 nm and dichroic long-pass filter at 504 nm and read with top optics, 200 flashes per well. Gain is adjusted using the only probe well with target mP of 35. Probe sequences are RMDSNIQGIENPGFEASPPAQGIPEAK for WT peptide and RMDSNIQGIEAAGFEASPPAQGIPEAK for NP→AA mutant peptide. Both peptides are N-terminal modified with 5-FAM-Ahx. Assays were performed in JS buffer (50 mM Hepes, pH 7.5, 100 mM NaCl, 1% glycerol, 1% Triton X-100, 1.5 mM MgCl2, and 5 mM EGTA) with peptide concentration 78.125 nM in total volume of 30 µL.

### Antibodies, chemicals, plasmids

Immunoblotting & immunofluorescence antibodies: α-VISTA (Cell Signaling #64953; R&D #MAB71261), α-EGFR (Cell Signaling #426*7*), α-phospho-EGFR Y1068 (Cell Signaling #3777), α-phospho-Akt S476 (Cell Signaling #4058), α-Akt (Cell Signaling #4685), α-phospho-ERK (Cell Signaling #9101), α-ERK (Cell Signaling #4695), α-PD-L1 (Cell Signaling #13684), α-actin (Invitrogen #MA515452), α-NUMB (Cell Signaling # 2756S), α-Rab11FIP1 (Cell Signaling #12849S), α-Rab11 (Cell Signaling #5589S), α-Rab7 (Cell Signaling #9367), α-caveolin (Cell Signaling #3267), α-EEA1 (Cell Signaling # 3288), α-LAMP1 (Cell Signaling #9091). Chemicals: TMR-dextran (Fisher Scientific #D1818), CellTiter-Blue (Promega #G8081). NUMB2-eGFP-M107K was a gift from Rolf Bjerkvig (Addgene plasmid # 37803). Human VISTA and VISTA-ΔC expression plasmids were constructed by ordering human VISTA (*VSIR)* sequence as a gBlock from IDT and performing Gibson assembly into the pLenti6.3/V5-DEST plasmid (Thermo Fisher), which was amplified with PCR primers (5’-GCAGAATTCCACCACACTGG, 5’-GGTAAGCCTATCCCTAACCCTCTCCT). miniTurboID sequence [57] was fused to VISTA and VISTA-ΔC by ordering gBlocks from IDT and performing Gibson assembly using PCR primers (5’-GGTAAGCCTATCCCTAACCCTCTCCT, 5’-GATGACCTCAAAGTTTGGAGAGTCAGGG for VISTA; 5’-GGTAAGCCTATCCCTAACCCTCTCCT, 5’-GTTAACGACCAGGAGCAGGATGAGG for VISTA-ΔC).

### Triple-negative breast cancer tissue microarray and immunohistochemistry

The tissue microarrays (TA459 and TA460) were constructed using a manual tissue arrayer (Beecher Instruments, Silver Spring, Maryland, United States) following previously described techniques [58] using 1.0 mm cores. Breast cancer samples were obtained from archived material at the Stanford University Medical Center Department of Pathology between 2000 and 2011 with IRB approval. The cores were taken from areas in the paraffin block that were representative of the diagnostic tissue based on examination of the original H&E slides. Tissue blocks from UTSW were obtained with IRB approval and sectioned onto glass slides in 6-8 samples per slide. Serial sections of 4uM were cut from tissue array blocks, baked at 60C for one hour, deparaffinized in xylene, and hydrated in a graded series of alcohol. Antigen retrieval was performed in SignalStain EDTA Unmasking Solution (Cell Signaling Technology, 14747) using a pressure cooker set to 95 °C for 3 minutes. Antibody staining was performed following the manufacturers protocol using the EnVision+ Dual Link System-HRP Rabbit/Mouse, DAB+ (DAKO, K4065) with the following changes: 1) addition of a 10-minute incubation with 3% hydrogen peroxide prior to the dual endogenous enzyme block and 2) blocking for 30 minutes with 3% goat serum at room temperature prior to primary antibody. The primary antibody VISTA D1L2G rabbit monoclonal antibody (Cell Signaling Technology, 64953) was diluted 1:50 in antibody diluent (DAKO, S0809) and incubated overnight in a humidified chamber at 4°C.

### Mouse studies

C57Bl/6j, Balb/cJ and NSG mice were purchased from Jackson Labs and housed in accordance with IACUC-approved animal protocols. For orthotopic mammary fat pad injections, mice were shaved or treated with Nair, then injections were performed with 15,000 – 50,000 cells per tumor for the 4T1 cell line or 300,000 – 500,000 cells per tumor for the HCC1806, MDA-MB-231 and E0771 cell lines into the mammary fat pad. Tumor growth was measured by digital calipers

### CRISPR Knock Out of VSIR and NUMB

To one well of a 96 well plate mix 95.76 μL Opti-MEM (Thermo Fisher Scientific, Catalog #31985062) with 0.24 μL lipofectamine RNAiMAX (Thermo Fisher Scientific, Catalog #13778030) and 12 μL of 6 μM of each sgRNA. The sgRNA sequences for VSIR are GACAUACAGGGAAUACGGCG (+71761997) and CUCUCUCUGAGCAGGUCCGG (−71762024), which were combined in a single transfection. The sgRNA sequence for NUMB is CUAUCGUCUGGUCAACUAUG (+73292867) and for NUMBL is UUCAUGGUGCCCGCCCCGUC (+40684555), which were combined in a single transfection. Then 120 μL (4000 cells) doxycycline-induced Cas9-expressing HCC1806 cells Opti-MEM media with 10% FBS to each well of a 96-well plate. After 24 h, 140 μL of the spent medium was removed and 100 μL of new growth medium was added. When cells were confluent, they were cloned by serial dilution (100 cells to 3 x 96-well plates) to isolate single cell clones, which were then expanded and screened for VISTA or NUMB protein levels by immunoblot. SgRNAs were purchased from Synthego, which contained an appended Synthego-modified EZ scaffold.

### TNBC tissue microarray analysis

VISTA immunohistochemistry scoring on tumor cells was performed by a blinded breast pathologist by the following algorithm: 0) negative: no VISTA stain present; 1+) indeterminate: cannot distinguish between lymphocytes versus tumor cells, blushed stain, single cells strongly stained, strong apical stain suspicious for non-specific; 2+) positive: clusters of strong stain or diffuse weak stain; 3+) strong positive: strong and diffuse membrane stain. PD1 expression by immunohistochemistry was scored as follows: the staining percentage of PDL1 for (iv) tumor and (v) stroma/lymphocytes by immunohistochemistry; and (vi) lymphocyte density by H&E. PDL1 stain was scored manually by a pathologist. PDL1 tumor staining percentage was averaged across both cores per patient. TIL density was averaged across the two cores. Only cases with interpretable scores for all scored variables and not lymph node metastases were included in downstream analysis (n=103). Welch’s two sample, unpaired, two-sided t-test with confidence level 0.95 was used to assess significance. In the UTSW cohort Ki-67 levels were extracted from clinical pathology reports.

### High-throughput Sequencing Analysis

RNA-seq and ATAC-seq datasets analyzed herein were previously deposited (GSE107119, GSE107121). The Protein Interactions Quantification (PIQ) footprinting software was applied using the NFKB1 motif from the JASPAR motif database. The pwmmatch.exact.r Rscript was run to create hits for the NFKB1 position-weight matrix in the hg19 genome. Then bam2rdata was run to convert the bam file to binary format. Then the pertf.r script then quantitated transcription factor binding scores based at each motif site for each bam file. Forward and reverse strand calls were combined, then only calls with purity score > 0.8 were retained. Peaks were converted to Granges and annotated with annotatePeakInBatch function of the ChIPpeakAnno package using EnsDb.Hsapiens.v75. Analysis was restricted to genes linked within 5 kb of a NFKB1 site. NFKB1 sites were restricted to those upstream of the TSS. The resulting gene list was used to filter the RNA-seq expression data and the selected genes were used in a volcano plot of differential gene expression +/- EGF. TCGA analysis was performed with the Gepia tool [59].

### Immunofluorescence analysis

CellProfiler software was applied to 63x pictures to identify nuclei from DAPI channel (pixel size range 15-45) with global threshold using two-classes Otsu method (smoothing factor=1.3488, threshold correction factor=1.0, lower bound =0, upper bound=1.0), then nuclei were expanded by 20 pixels. Lysosomes or vesicles were identified based on size (6-18 pixels) with global threshold using two-class Otsu method (smoothing factor=0, correction factor=1.0, lower bound=0, upper bound=1.0). Feature sizes were then measured in pixels and feature intensity was derived. RelateObjects command was used to assign vesicle features to nuclei based on the expanded nuclei maps to calculate per-parent statistics for each child. Prism was then used for further statistical analysis. Manders’ colocalization coefficient was calculated with JaCoP.

