## Supplementary Materials for "VISTA-induced tumor suppression by a four amino acid intracellular motif"

^*^Corresponding Authors:

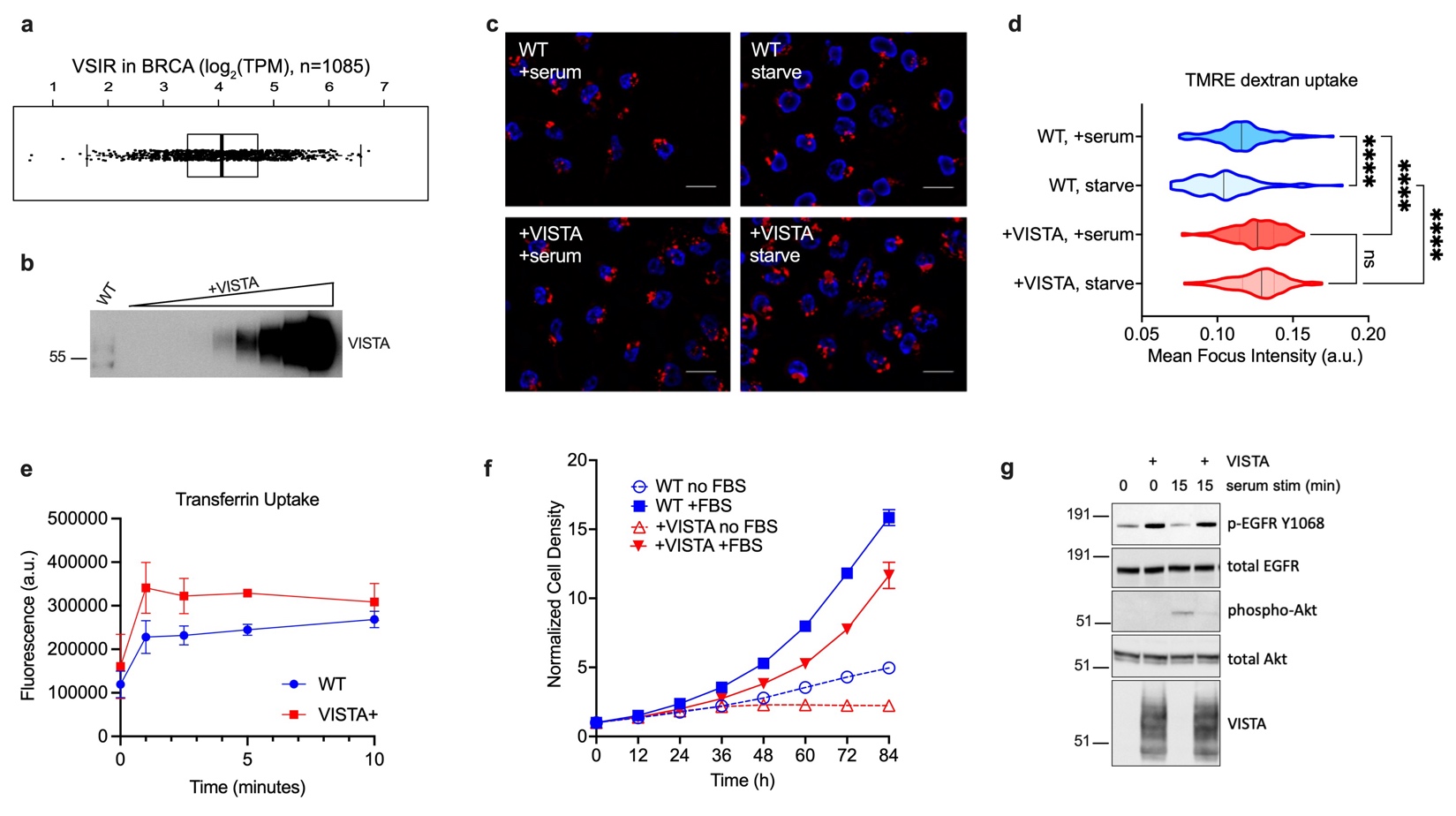
**Supplementary Fig. S1:** VISTA expression, macropinocytosis, transferrin uptake and serum stimulation in VISTA+ cells. **a.** VSIR transcript levels in the TCGA BRCA dataset. **b.** Quantitative immunoblots for VISTA levels in serum starved WT HCC1806 cells versus serial dilutions of VISTA+ cells. **c.** HCC1806 cells were grown +/- serum for 24 h then incubated with tetramethylrhodamine-dextran and imaged by fluorescence microscopy. Scale bar 25 μm. **d.** Quantitation of mean fluorescence intensity per foci per cell from b. **** p < 0.0001 by Kruskal-Wallis. N = 552 cells total **e.** Transferrin receptor endocytosis assay was performed in HCC1806 cells. mean +/- SD, N = 5 replicates per group. **f.** HCC1806 +/- VISTA were grown in serum-free or serum stimulated conditions and growth density was measured by Incucyte. FBS = fetal bovine serum. **g.** HCC1806 cells +/- VISTA were serum starved for 48 hours then stimulated with serum for 15 minutes, then proteins were analyzed by immunoblot.

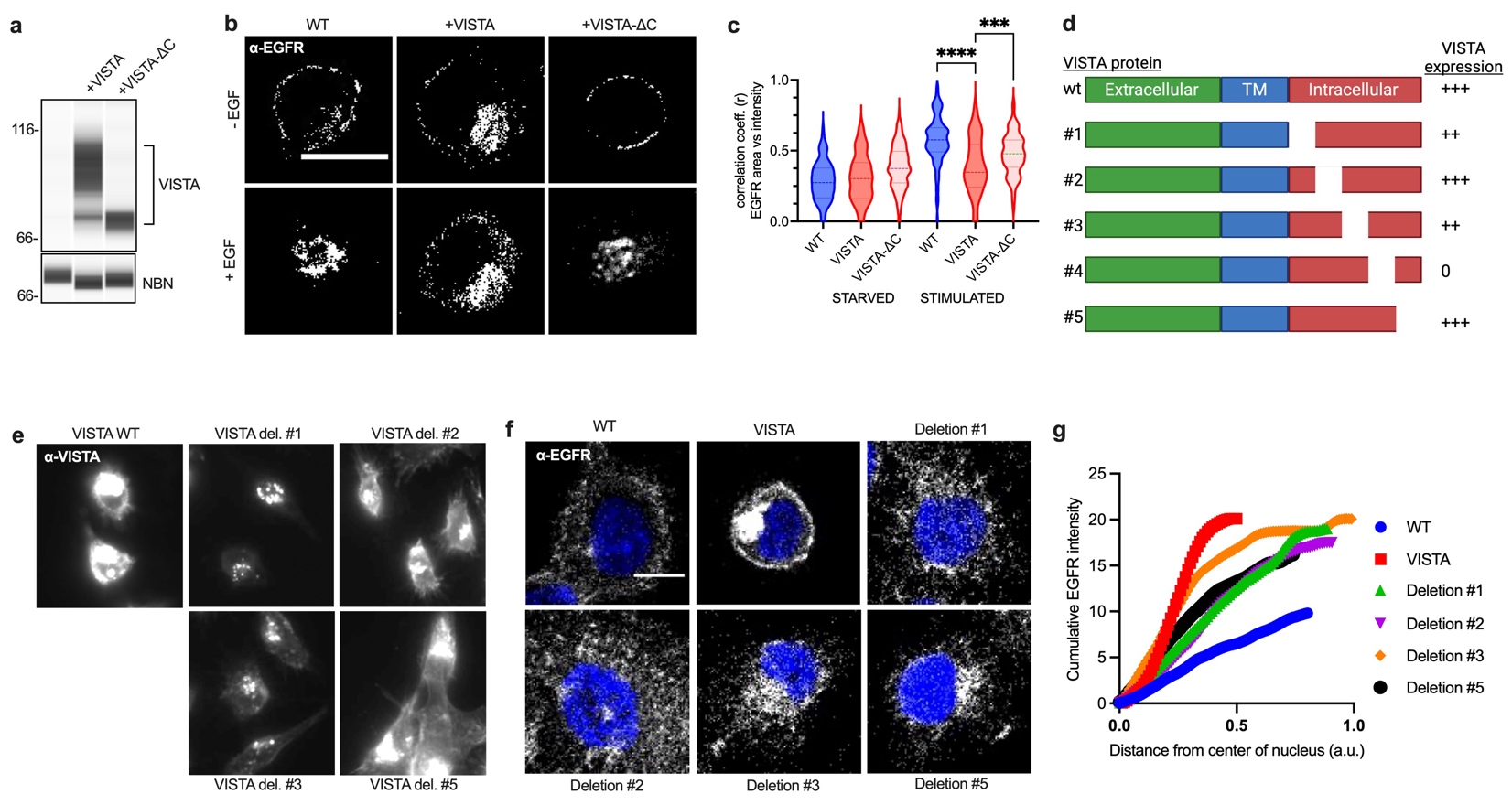
**Supplementary Fig. S2:** VISTA-CTD controls EGFR localization. **a**. Microcapillary immunoassay for protein levels in HCC1806 cells (WT, +VISTA, +VISTA-ΔC). **b.** HCC1806 cells (WT, +VISTA, +VISTA-ΔC) were starved of serum for 48 hours, then stimulated with EGF for 30 minutes and EGFR was detected by confocal microscopy. Scale bar = 20 μm. **c.** Quantification of EGFR localization changes from b. **d.** Schematic of VISTA CTD deletions constructed and their relative expression in cells as detected by immunofluorescence microscopy. **e.** Immunofluorescence images of VISTA WT and deletion mutants expressed in HCC1806 cells and detected by antibodies towards the VISTA extracellular domain. **f.** Confocal microscopy images of EGFR localization in HCC1806 cells expressing WT VISTA or deletion mutants. **g.** Quantitation of EGFR intensity in a vector from the center of the nucleus through the highest EGFR density towards the cell membrane for images in e (n=10 cells per sample).

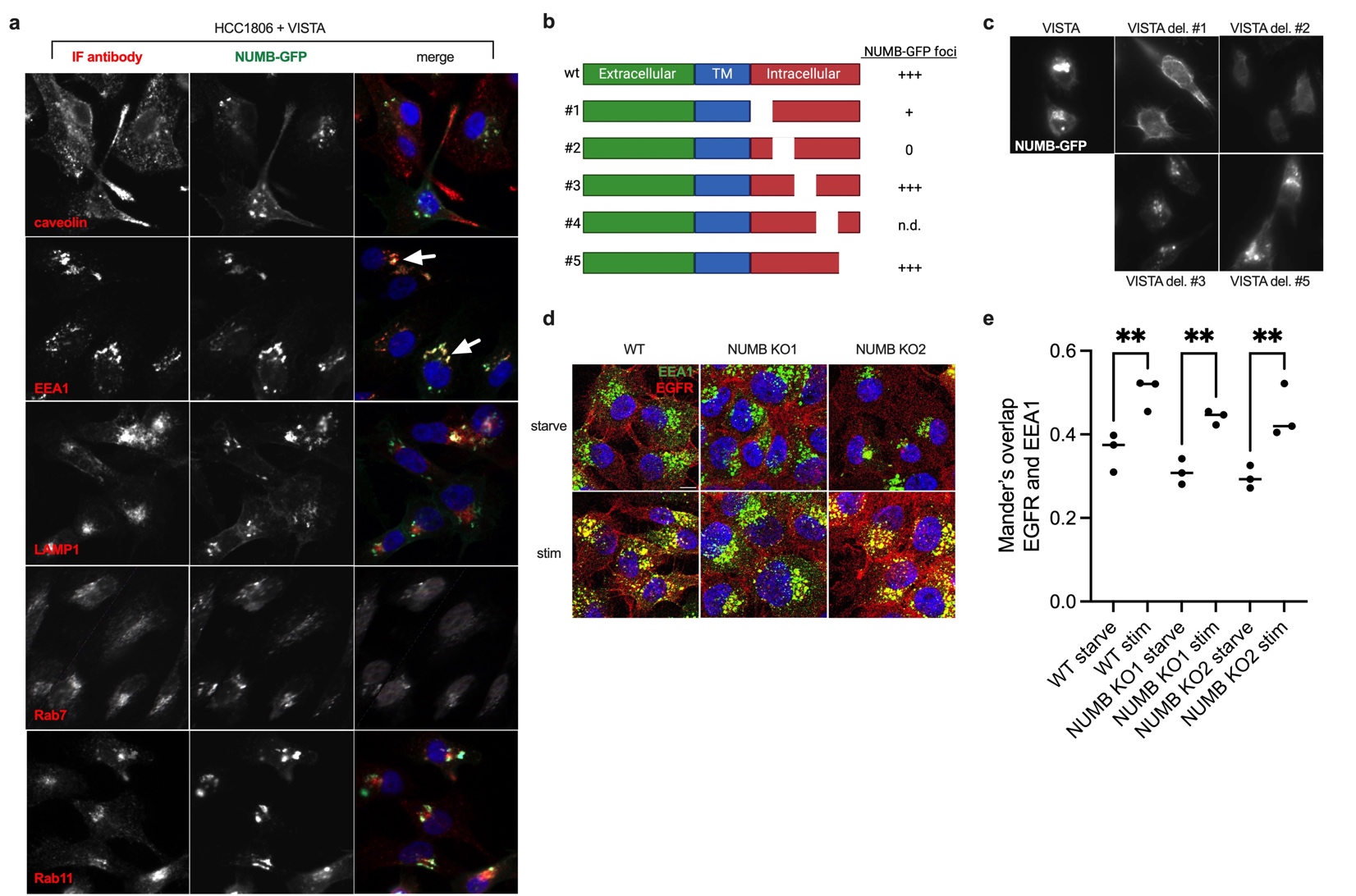
**Supplementary Fig. S3:** VISTA’s biochemical and genetic interactions with NUMB. **a.** NUMB-GFP fusion protein was expressed in HCC1806 VISTA+ cells and immunofluorescence was performed for indicated antibodies (red). Scale bar is 20 μm, arrows show co-localization. **b.** Schematic of VISTA-CTD deletion mutants with the formation of NUMB-GFP foci scored. n.d. = not determined **c.** Representative fluorescence microscopy images of NUMB-GFP localization in HCC1806 cells that express VISTA-CTD mutants. **d.** Confocal imaging of EGFR (red) and EEA1 (green) localization in WT and NUMB KO HCC1806 cell lines, starved 48 h or stimulated 30 min with serum + EGF. Scale bar = 20 μm. **e.** Quantification of EGFR and EEA1 overlap from **d.** by Mander’s. Each point represents an independent image. ** p-value < 0.01 by ANOVA with Sidak’s test.
